# Prenatal exposure to the probiotic *Lactococcus lactis* decreases anxiety-like behavior and modulates cortical cytoarchitecture in a sex specific manner

**DOI:** 10.1101/780072

**Authors:** Natalia Surzenko, Eneda Pjetri, Carolyn A. Munson, Walter B. Friday, Jonas Hauser, Ellen S. Mitchell

## Abstract

Development of the cerebral cortex may be influenced by the composition of the maternal gut microbiota. To test this possibility, we administered probiotic *Lactococcus lactis* in the drinking water to mouse dams from day 10.5 of gestation and until pups reached postnatal day 1 (P1). Pups were assessed in a battery of behavioral tests starting at 10 weeks old. We found that females, but not males, exposed to probiotic during prenatal development spent more time in the center of the open field and also displayed decreased freezing time in cue associated learning, compared to controls. Furthermore, we found that probiotic exposure changes the densities of cortical neurons and increases the density of blood vessels in the cortical plate of P1 pups. Sex-specific differences were observed in the numbers of mitotic neural progenitor cells, which were increased in probiotic exposed female pups. In addition, we found that probiotics treatment throughout pregnancy significantly increased plasma oxytocin levels in mouse dams, but not in the offspring. These results suggest that exposure of naïve, unstressed dams to probiotic may exert sex-specific long-term effects on cortical development and anxiety related behavior in the offspring.

## Introduction

Neocortical development is influenced by a variety of environmental factors, including maternal nutrition, stress and exposure to pathogens. In terms of cortical patterning and growth, the impact of essential micronutrients, such as choline, has been well-characterized; yet less is known on how food-derived microbiota, e.g. probiotics, may be altering cortical development at different stages ^[1]^. While the fetal gut is sterile before birth, it is increasingly clear that factors from the maternal microbiota can induce specific patterns of gene expression in the fetal gut, as well as in the brain ^[1–3]^. After birth, microbial colonization of the infant gastrointestinal tract has been shown to have widespread influence on brain development and behavior ^[4]^. For example, mice whose gut microbiota were depleted via antibiotics or who were raised in a germ-free facility demonstrate exaggerated responses to stress and social stimuli, and display neurochemical and brain structural abnormalities ^[5, 6]^. Recently, neonatal exposure to probiotics, such as *Bifidus longum* and *Lactobacillus rhamnosous*, have been shown to reverse maladaptive learning behaviors in innately anxious mice ^[7]^. Probiotics have also alleviated behavioral effects in other models of anxiety disorders, such as those induced by maternal separation, social defeat or other types of early life stress ^[8–10]^. Mounting data on brain-gut axis has revealed several pathways where probiotic treatment can affect microbiota populations, brain signaling, and subsequently anxiety in mouse models of disease. However, there is less consensus on the behavioral effects of probiotics in naïve, wild type mice which have not been exposed to stress paradigms.

Neurochemical investigations have revealed that probiotics treatment reduced inflammatory cytokines and stress-related hormones, which are often chronically activated in anxiety disorders. For instance, modulation of the vagus nerve has been implicated in behavioral effects of probiotics, since vagal resection blocks the anti-inflammatory, anxiolytic activity of probiotics ^[11]^. Additionally, probiotics may be affecting brain development by improving maternal health and immune-mediated stress responses. Exposure to probiotics, such as *Lactobacillus reuteri*, can increase plasma oxytocin ^[12]^, potentially enhancing nurture instincts and pup handling in treated dams. Studies such as these implicate changes in neuroendocrine signaling brought on by plasma metabolites derived from the probiotics themselves or from specific commensal microbiota interacting with the probiotics. In fetuses with rudimentary microbiomes, metabolites from maternal microbiome access the fetal blood stream via placental transference. Yet, there is a dearth of knowledge on how probiotic exposure in utero, i.e. before postnatal colonization, affects structural brain development in wild type, naïve neonates. In the present study we investigated whether neonatal exposure to the probiotic *Lactococcus lactis* (*L. lactis*) induces long-lasting changes in cortical layer structure. We found that the densities of distinct types of cortical neurons were changed by probiotics exposure and associated with increased proliferation of cortical neural progenitor cells. Additionally, we measured anxiety and emotional learning capacity in probiotics-exposed mice and found that maternal L. lactis exposure ameliorates anxiety-related behavior in the offspring in a primarily sex-specific manner.

## Materials and Methods

### Animals

All experiments were performed at the David H. Murdock Research Institute Center for Laboratory Animal Science facilities in accordance with the standards of the U.S. National Institutes of Health Guide for Care and Use of Laboratory Animals and were approved by the Institutional Animal Care and Use Committee at this facility. *Nestin-CFPnuc* transgenic mice were generously provided by Dr. Grigori Enikolopov (Cold Spring Harbor Laboratory, Cold Spring Harbor, NY, USA) ^[25]^ and were maintained on a mixed C57BL/6J and C57BL6/N background (97% C57BL/6J).

Mice were habituated to modified AIN93G diet (#103186, Dyets Inc., Bethlehem, PA) for at least two weeks before breeding and until the end of the experiments. This diet contains 1.4 g choline chloride/kg diet, which meets the mouse requirements for methionine and choline (John and Bell 1976, Council 1995).

Pregnant dams were randomly divided in two groups: one group was given the probiotic *Lactococcus lactis* (*L. lactis*) in drinking water (5×10^5 CFU/ml) from gestational day (GD) 10.5 to postnatal day 1 while the other group (Control) received drinking water only. The drinking water was refreshed daily. Probiotic CFU was confirmed to remain unchanged after study completion.

### Behavioral tests

Mice were tested starting at 10-13 weeks in the behavioral battery for locomotor activity and anxiety-like behavior in open field, exploratory behavior in object investigation, anxiety-like behavior in the light-dark box test and for learning and memory in fear conditioning test ^[26]^. In the control group, nine male and ten female mice were tested, while in the *L.lactis* exposed group, ten males and 12 females were tested. The mice were littermates from multiple litters. The mice were kept on a 12:12 hr light-dark schedule and were tested in the light phase of the day. Male and female mice were tested on separate days and were given at least 48 hr between the tests.

### Open field test

This test is a widely used test to assess locomotor activity and as a measure of anxiety-like behavior in a novel, stressful environment ^[27]^. The open field was a circular arena with diameter 80 cm and height 35 cm that was evenly illuminated with white light. The mice were allowed to explore for 5 min while the test was recorded and analyzed with an automated tracking system, Ethovision XT® (Noldus Information Technologies, The Netherlands). We measured total distance moved, velocity, latency to center zone, and total duration in center zone.

### Object investigation test

In this test, we measured the exploratory activity of a mouse towards a novel non-aversive object placed in the center of the arena after they are familiar with the environment ^[28, 29]^. The mice were placed facing the wall in the same open field arena and allowed to habituate for 2 min. Then, the novel object is introduced in the center of the arena and the mice are allowed to explore it freely for 3 min. The total time investigating the object was scored manually.

### Light-dark box

The light-dark exploration test is used to measure anxiety-like behavior and the conflict between rodents’ exploratory behavior and aversion to open and brightly illuminated areas ^[30]^.

The test box (44 × 21 × 21 cm; l × w × h) is divided into two unequal compartments by a partition (1 cm) with a small aperture (5 × 7 cm) in the center by the floor level, allowing the mice to move freely between the chambers. The dark chamber, 1/3 of the box, is black and covered with a lid. The light chamber is white and brightly illuminated. The test mouse was placed into the dark chamber facing the end wall and allowed to explore for 5-min. Ethovision XT® was used to record locomotor activity in the light zone. Transition between chambers, when all four paws are placed in one chamber, was scored manually. Duration in each chamber and the latency to go to the light chamber was also recorded.

### Contextual and cued fear conditioning test

Contextual and cued fear conditioning was performed using a conditioned fear paradigm over 3-days using Near-Infrared image tracking system (MED Associates, Burlington, VT) ^[31]^. On the first day, after an initial 2-min exploration time (background activity), mice were exposed to a 30-s tone (85 dB, 2800Hz), followed by a 2-s scrambled foot shock (0.75mA) (CS1) under white light conditions. Mice received 3 additional tone-shock pairings (CS2 and CS3), with 80-s between the stimuli pairings, totaling a 9.2-min session. The response to the shock was measured with automatic assessment of the levels of freezing (immobility, except for breathing) using the Video Freeze (MED Associates Inc.) software. All mice learned the association between the tone and the shock, and the response to the tone increased with each pairing.

### Immunohistochemistry

P1 brains, representing pups from 4-5 independent litters, were fixed in 4% PFA, cryopreserved through incubation in 10%-20%-30% sucrose/1 X PBS gradient over 72 hours at 4°C, mounted in O.C.T. compound and stored at −20°C. Brains were cryosectioned coronally at 20 μm over a series of 7 slides, such that each slide contained representative non-consecutive brain sections. Slides were re-hydrated in 1X PBS and incubated in blocking solution containing 2% goat serum/0.01% Triton-X in 1X PBS for 1 hour at room temperature. Antibodies were dissolved in blocking solution and applied over night at 4°C as follows: anti-PECAM1 (rat, 1:50; Thermo Fisher Scientific); anti-PH3 (rabbit, 1:1000; Millipore-Sigma) anti-Satb2 (mouse, 1:250; Abcam); anti-Tbr1 (rabbit, 1:250; Abcam). DAPI (1:4000; Millipore-Sigma) was used to detect cell nuclei. The following secondary antibodies were used: goat anti-rabbit CY3 (1:250; Jackson Immunoresearch); goat anti-mouse Alexa488 (1:1000; Jackson Immunoresearch); goat anti-rat CY3 (1:250; Jackson Immunoresearch).

### Image acquisition and cell density analyses

Image z-stacks were acquired using Zeiss LSM710 laser scanning confocal microscope and 20X objective. Obtained stacks spanned 12-14 μm and contained 5-7 optical slices. Brains were analyzed at the level of the presumptive visual cortex. ImageJ (NIH) software was used in manual and automated cell density analyses. Tbr1-expressing cells were manually counted in 60 × 60 μm regions of the lower cortical plate. Satb2-expressing cells were manually counted in 60 × 60 μm regions of the upper cortical plate. PECAM1-expressing blood vessels were manually counted in 100 × 100 μm regions of the upper cortical plate; fluorescence intensity and blood vessel areas were measured using ImageJ automated particle analysis module. PH3-expressing cells in the ventricular zone were manually counted on images collected at 10X magnification.

### Oxytocin measurements

Dams were administered probiotics in water, or water alone (control) starting at day 10.5 of pregnancy and plasma was collected 1 day after birth. Plasma oxytocin was extracted and measured according to the protocol supplied by the manufacturer of the ELISA kit (Enzo Life Sciences; ADI-900-153A-0001). Measurements were conducted using BioTek Synergy2 plate reader.

### Statistical analyses

Statistical analyses were performed with IBM SPSS Statistics (behavioral assays) and Graphpad (Prism) software. Data were checked for outliers using Grubbs test, normality using the Shapiro-Wilk test of normality and, for homogeneity of variances using Levene’s test. To account for sex effects, two-way ANOVA analysis, with treatment (control and probiotic) and sex (male and female) as factors, was used to analyze the data. When there was a significant sex effect, a separate analysis by sex was conducted. If the data were not normally distributed, non-parametric Mann-Whitney test was used. In the fear conditioning experiment, repeated-measures ANOVA was also used to analyze the freezing behavior during the tests. Results are expressed as mean ± SEM unless otherwise specified.

## Results

### Maternal probiotic supplementation reduces anxiety-like offspring behavior in light-dark box test

To evaluate whether prenatal exposure to probiotics affects anxiety-like behavior, mice were tested in the open field and light-dark box. In the light-dark box test, even though the *L. lactis*-exposed mice spent 10% more time in the light zone than the controls, the difference didn’t reach significance with two-way ANOVA analysis for either simple main effects or interaction between these effects (p > 0.05, Figure 1A). However, there was a significant interaction between the effects of sex and treatment on activity levels in the light zone (F(1,36) = 4.54, p = 0.04, Figure 1B). Main effects analysis showed a trend for significance for the effects of in utero *L. lactis* exposure (F(1,36) = 3.817, p = 0.059), but there were no differences between males and females (p = 0.927).

**Figure 1.**
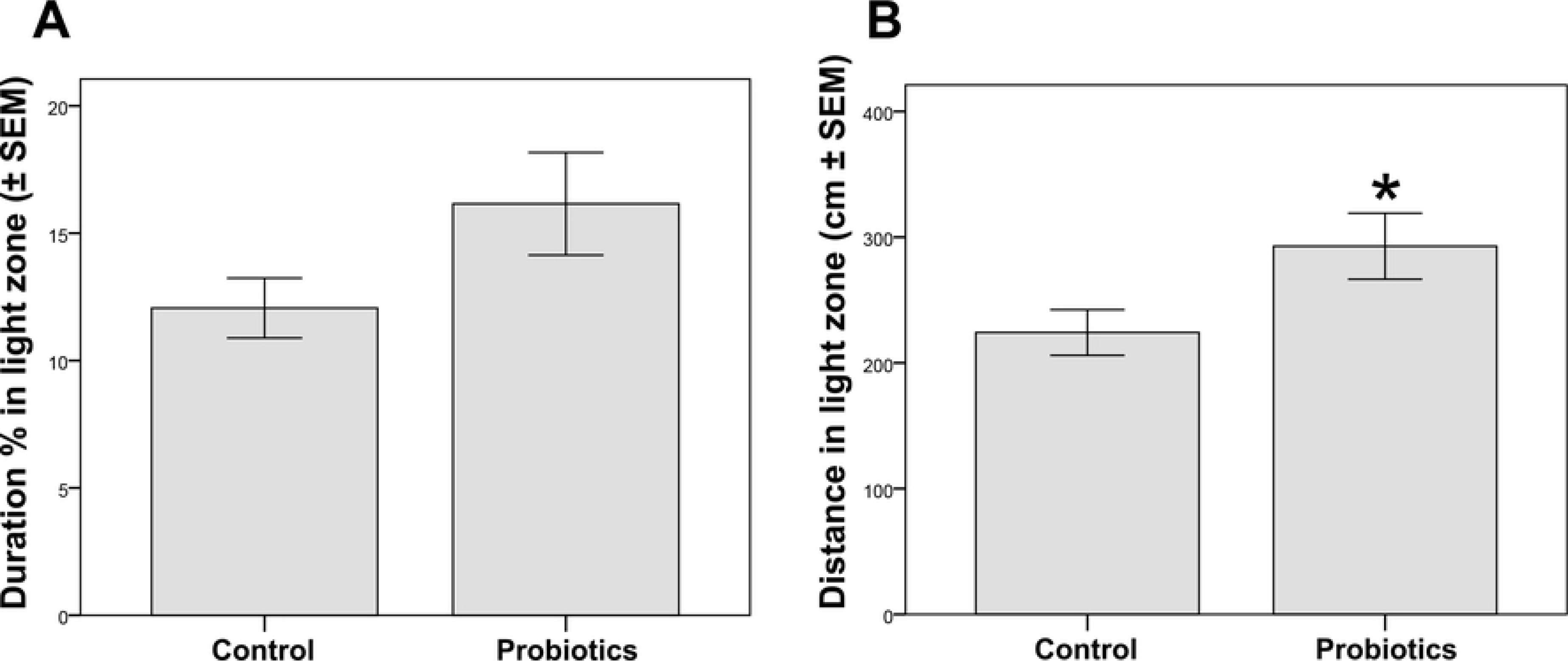
Maternal probiotic supplementation reduces anxiety-like offspring behavior in light-dark box test. (**A**) Probiotic-exposed mice (male n = 9 and female n = 12) spent 10% more time than the controls ((male n = 9 and female n = 10) in the light zone, however this didn’t reach significance. (**B**) There was a significant interaction between the effects of sex and treatment on activity levels in the light zone (p = 0.04).

In the open field, we observed a decrease in the latency to enter the center zone in *L. lactis*-exposed mice. However, this decrease didn’t reach significance (Control 49.09 ± 15.7s and *L. lactis* 17.73 ± 3.9s, p = 0.060, Supplemental Figure S1A) and there were no sex differences (p > 0.05). The activity levels were similar between the groups (Control 2547 ± 112 cm and *L. lactis* 2795 ± 103 cm, p > 0.05) and there were no sex differences (p > 0.05). We also looked at the time spent in the center as another measure of anxiety-like behavior. We didn’t find an effect of *L. lactis* treatment (Control 6.84 ± 1.63 cm and *L. lactis* 7.04 ± 1.38 cm, p > 0.05), but there was a strong main effect of sex, with female mice spending significantly more time in the center than male mice (F(1,37) = 100, p < 0.001, Figure S1B).

In the novel object exploration test, there was no difference in the latency to approach the object (Control 45.56 ± 9.53 s and *L. lactis* 29.98 ± 7.22 s, p > 0.05 Figure S2A), and there were no main effects of sex. Time spent investigating the object was not different between the groups (Control 8.94 ± 2.15 s and *L. lactis* 11.23 ± 2.86 s, p > 0.05 Figure S2B).

### Maternal probiotic supplementation differentially affects female mice in fear conditioning test

Treatment with probiotic did not affect conditioning, when analyzed with repeated measures ANOVA (p > 0.05). There was a significant interaction between sex and group (F(1.34) = 4.257, p = 0.047) with control female mice showing higher levels of freezing than males, however, the main effect of sex did not reach significance (p = 0.06 Fig 2A).

**Figure 2.**
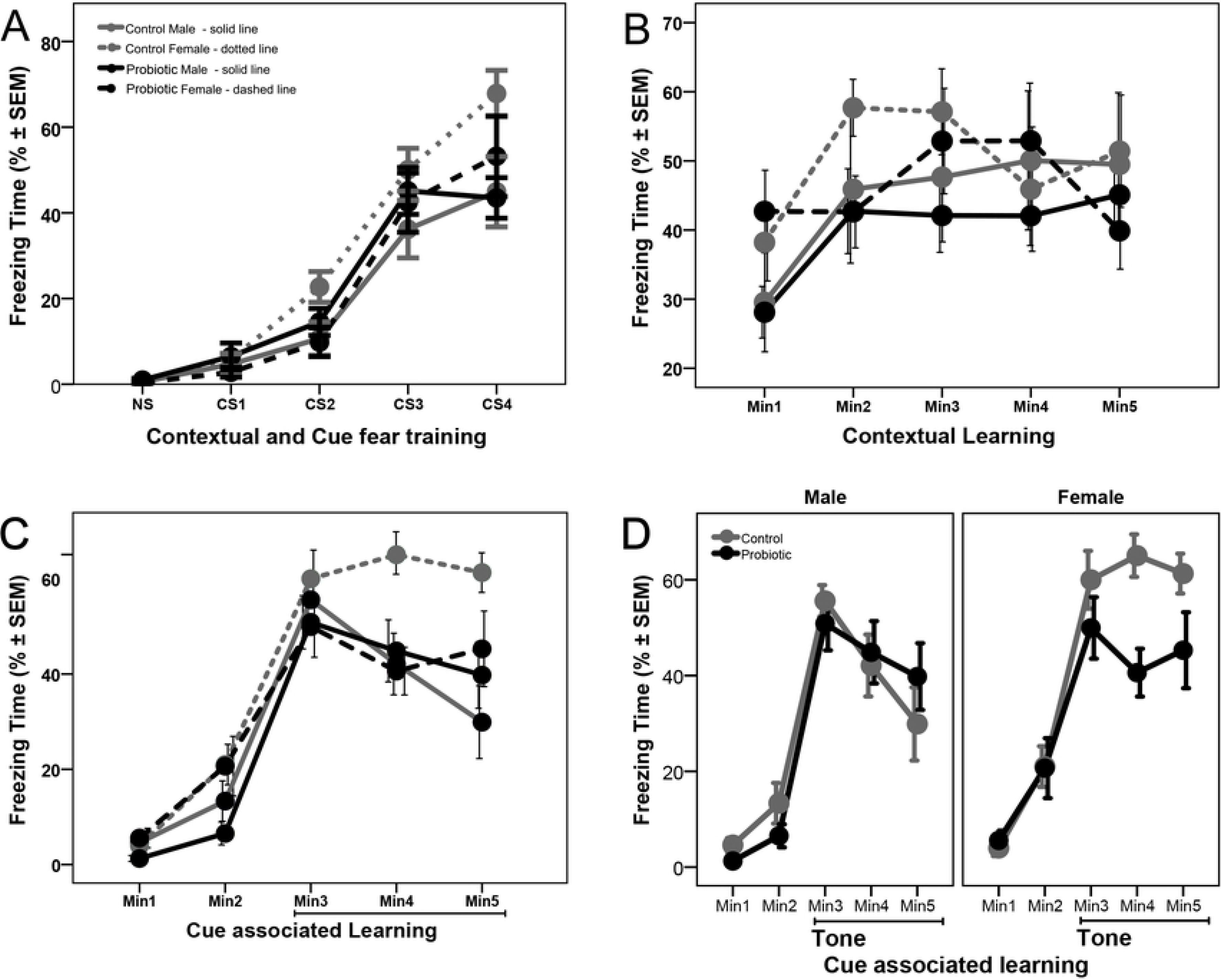
Maternal probiotic supplementation differentially affects female mice in fear conditioning test. (**A**) In utero exposure to probiotics did not affect conditioning of the mice. Control female mice (n = 10) showed higher levels of freezing than males (n = 8), however, the main effect of sex did not reach significance (p = 0.06). (**B**) Probiotic exposure did not have an effect on the contextual memory of mice. Though female mice (n = 10) showed higher levels of freezing than male mice (n = 10), separate sex analysis showed no treatment effect. (**C**) Probiotic exposure did not affect the first minute response to tone, but there is a significant treatment effect in female mice, shown in detail in panel (**D**).

On the following day, mice were evaluated for their context-dependent learning. Mice were placed back into the original test chamber for a 5-min session, this time with no tone or foot shock. Probiotic treatment did not have an effect on the first minute response (p > 0.05) and there was no interaction between group and sex with univariate ANOVA (p > 0.05). However, there was a significant sex effect (F(1.34) = 4.287, p = 0.046) with female mice showing higher levels of freezing. Separate analysis in female and male mice showed no treatment effect (p > 0.05, Figure S3A). Repeated measures analysis, for the 5 minutes of the session, showed no differences between the groups and also no effect of sex (p > 0.05, Figure 2B). Analysis of average freezing time during the 5-min session showed no effect of treatment or sex (p > 0.05, Figure S3B).

On the third day of testing, associative learning to the tone cue was evaluated. The conditioning chambers were modified by turning off the white light and keeping only Near-Infrared light, by modifying the chamber using a black Plexiglas insert in an A-shape to change the wall and another insert to change the floor surface, and, by adding a novel odor (vanilla flavoring). Mice were placed in the modified chamber and allowed to explore it for a final 5-min session. After 2-min, the acoustic stimulus was presented continuously for a 3-min period. Compared to control, probiotic treatment did not have an effect on the first minute response to tone (p > 0.05) and there was no interaction between group and sex, or a sex effect with univariate ANOVA (p > 0.05, Figure S4A). Repeated measures analysis for the 3 minutes of tone presentation showed no differences between the groups (p > 0.05) and neither the interaction between groups and sex, nor sex effect, reached statistical significance (p = 0.067 and p = 0.064 respectively, Figure 2C). However, separate analysis in female and male mice showed a significant treatment effect with repeated measures analysis for the 3 minutes of tone presentation in female mice (F(1.18) = 5.700, p = 0.028, Figure 2D) but not in male mice (p > 0.05). Treatment with probiotic reduced the freezing time in female mice in the second and third minutes of tone presentation, while the female mice in the control group maintained similar freezing levels. Overall, the above results suggest that probiotics are predominantly affecting female behavior after *in utero* exposure.

### *L. lactis* exposure during pregnancy induces an increase in maternal oxytocin levels

Oxytocin is a known hormonal mediator of maternal behavior ^[13]^. To examine whether *L. lactis* exposure during pregnancy may increase maternal oxytocin levels, we measured oxytocin in plasma of treated and control dams 1 day after birth (Figure 3). We find that oxytocin levels are significantly increased in plasma of *L. lactis*-treated dams compared to control dams (Figure 3A; p = 0.0008 by Student’s t-test). Oxytocin levels were not increased in plasma of pups at P28 (Figure 3B). These results suggest that probiotics exposure during pregnancy may modulate the levels of maternal oxytocin.

**Figure 3.**
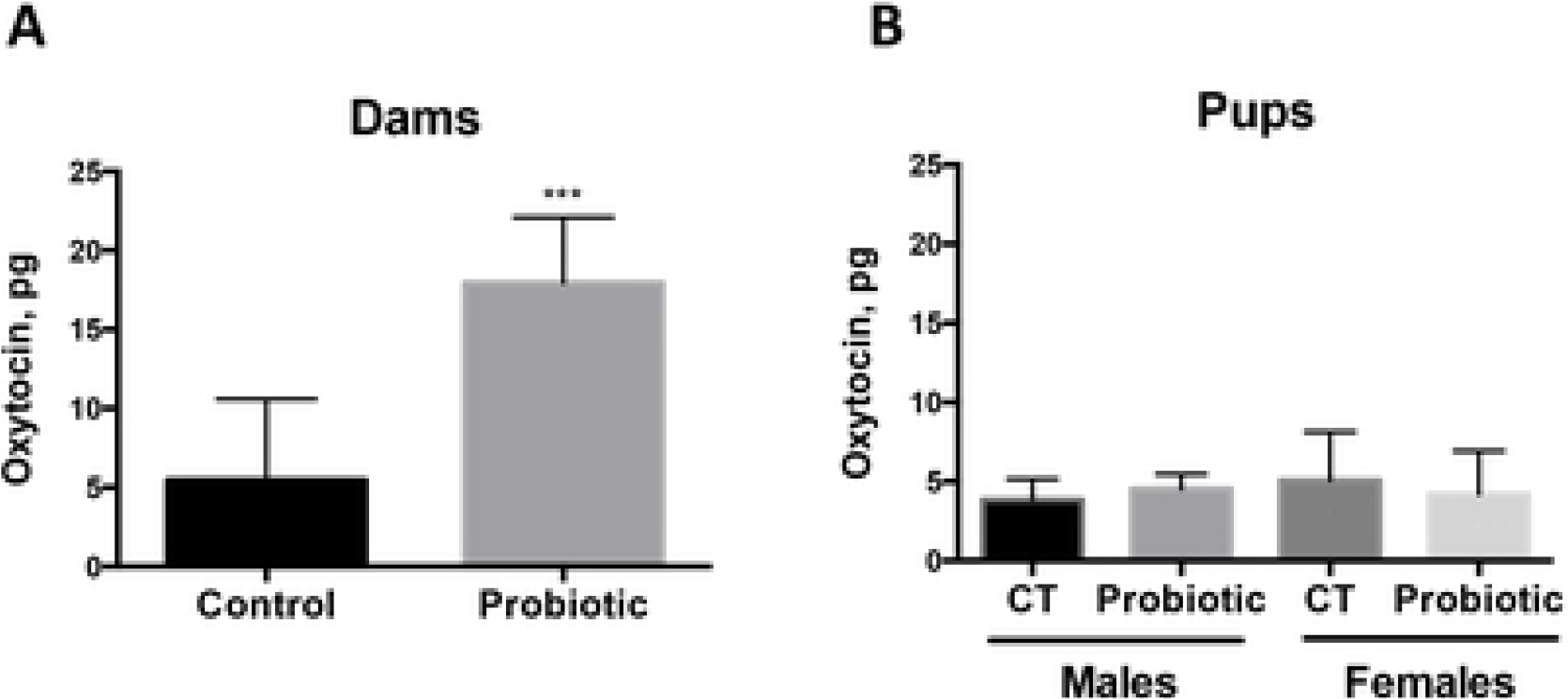
Increase in plasma oxytocin levels in probiotics-treated dams. (**A**) Oxytocin levels are increased in the plasma of dams treated with *L. lactis* between day 0.5 of pregnancy and 1 day after birth (n = 5) compared to Control dams (n = 8). P = 0.0008 by Student’s t-test. (**B**) Plasma oxytocin levels remain unchanged in 3-month old male and female pups exposed to *L. lactis* between E0.5 and P1 (n = 5-6 per sex/group).

### Maternal supplementation with *L. lactis* modulates development of cortical vasculature in the pups

Maternal gut microbiota is known to influence blood vessel permeability in the developing embryo ^[14]^. In this study, we examined whether the process of blood vessel formation during cortical development is affected by maternal exposure to *L. lactis*. To this end, we evaluated the expression of PECAM1 (CD31), an endothelial cell marker, in the developing cortices of *L. lactis*-exposed compared to control P1 pups. We find that blood vessel number in the cortical plate is increased in probiotics-exposed male (p = 0.0007 by Student’s t-test) and female (p = 0.0037 by Student’s t-test) pups compared to control pups (Figure 4A-C). Furthermore, expression levels of PECAM1, measured by fluorescence intensity, is increased in *L. lactis*-exposed pups (Figure 4B, E; p = 0.0002 by Student’s t-test). Finally, analysis of blood vessel areas reveals an increase in the average blood vessel area in probiotics-exposed compared to control pups (Figure 4F; p = 0.0006). Together, these results demonstrate that maternal exposure to probiotics modulates formation of the vasculature in the developing embryonic brain.

**Figure 4.**
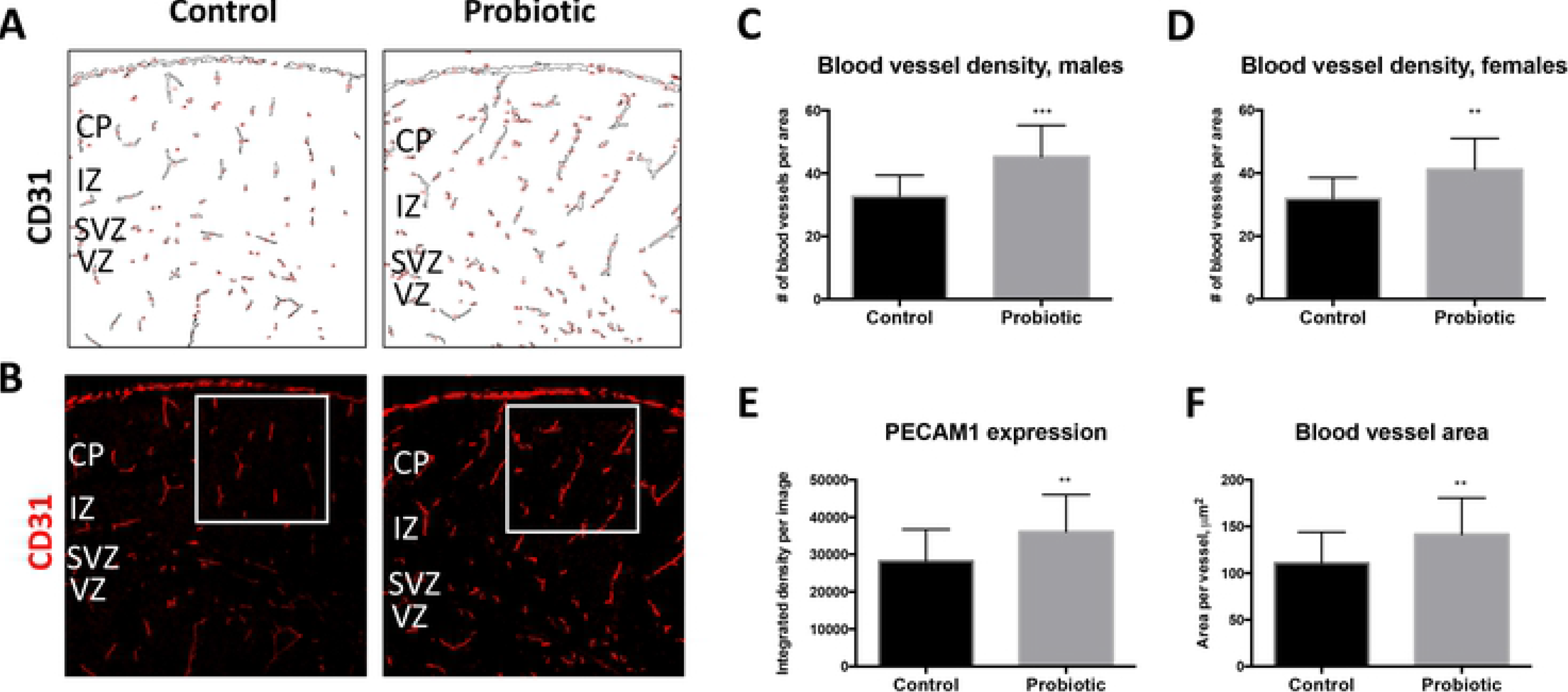
Maternal exposure to probiotics modulates formation of cortical vasculature in the pups. (**A**) Blood vessel outlines based on CD31 expression demonstrate increase in the density of cortical blood vessels in probiotic-exposed P1 pups compared to control pups. Counting masks are shown in red. (**B**) Immunofluorescent images of CD31 staining show increased staining intensity in P1 *L. lactis*-exposed compared to control cortices. White squares designate areas of blood vessel density analyses. (**C-D**) Increase in blood vessel density within the cortical wall is detected in both male (**C**; p = 0.0007 by Students t-test; control n = 18; *L. lactis* n = 9) and female (**D**; p = 0.0037 by Two-tailed Student’s t-test; control n = 15; *L. lactis* n = 20) *L. lactis*-exposed versus control P1 pups. (**E**) Average expression levels of CD31 per blood vessel is increased in *L. lactis*-exposed P1 pup cortices (p = 0.0045 by Two-tailed Student’s t-test; control n = 33; *L. lactis* n = 18). (**F**) Average blood vessel area is increased in P1 probiotic-exposed compared to control pups (p = 0.0045 by Two-tailed Student’s t-test; control n = 33; *L. lactis* n = 18). Scale bar in (**B**): 80 μm. *CP* - cortical plate; *VZ* - ventricular zone.

### *In utero* exposure to *L. lactis* leads to structural changes in pyramidal neuronal cell layer organization of the cerebral cortex

Mouse cortical neurogenesis begins around embryonic day 11.5 (E11.5) and continues throughout gestation while cortical gliogenesis peaks during postnatal development ^[15]^. Cortical pyramidal cell layers are generated in the inside-out manner, such that lower layer neurons (layers VI and V) are produced early in development, followed by the upper layer neurons (layers VI-II) ^[16]^. To determine whether maternal supplementation with *L. lactis* from E10.5 through postnatal day 1 (P1) affects cellular composition of the developing cortex we examined the expression of the cortical lower layers marker, Tbr1 (layer VI), and upper layers marker, Satb2 (layers II-V), at P1. We find that the density of Tbr1-expressing neurons in *L. lactis*-exposed pups is increased (Figure 5A-D; p < 0.0001 and p = 0.05, respectively, by Student’s t-test). However, cortical layer VI thickness, as determined by the thickness of Tbr1-positive cell layer, is not statistically significant between groups in either male or female pups (Figure 5A (bracket), E-F). Similarly, the density of Satb2-expressing layer II-IV neurons is also increased, albeit only in male probiotics-exposed pups (Figure 5B, G; p = 0.0006 by Student’s t-test) (Figure 5B, H).

**Figure 5.**
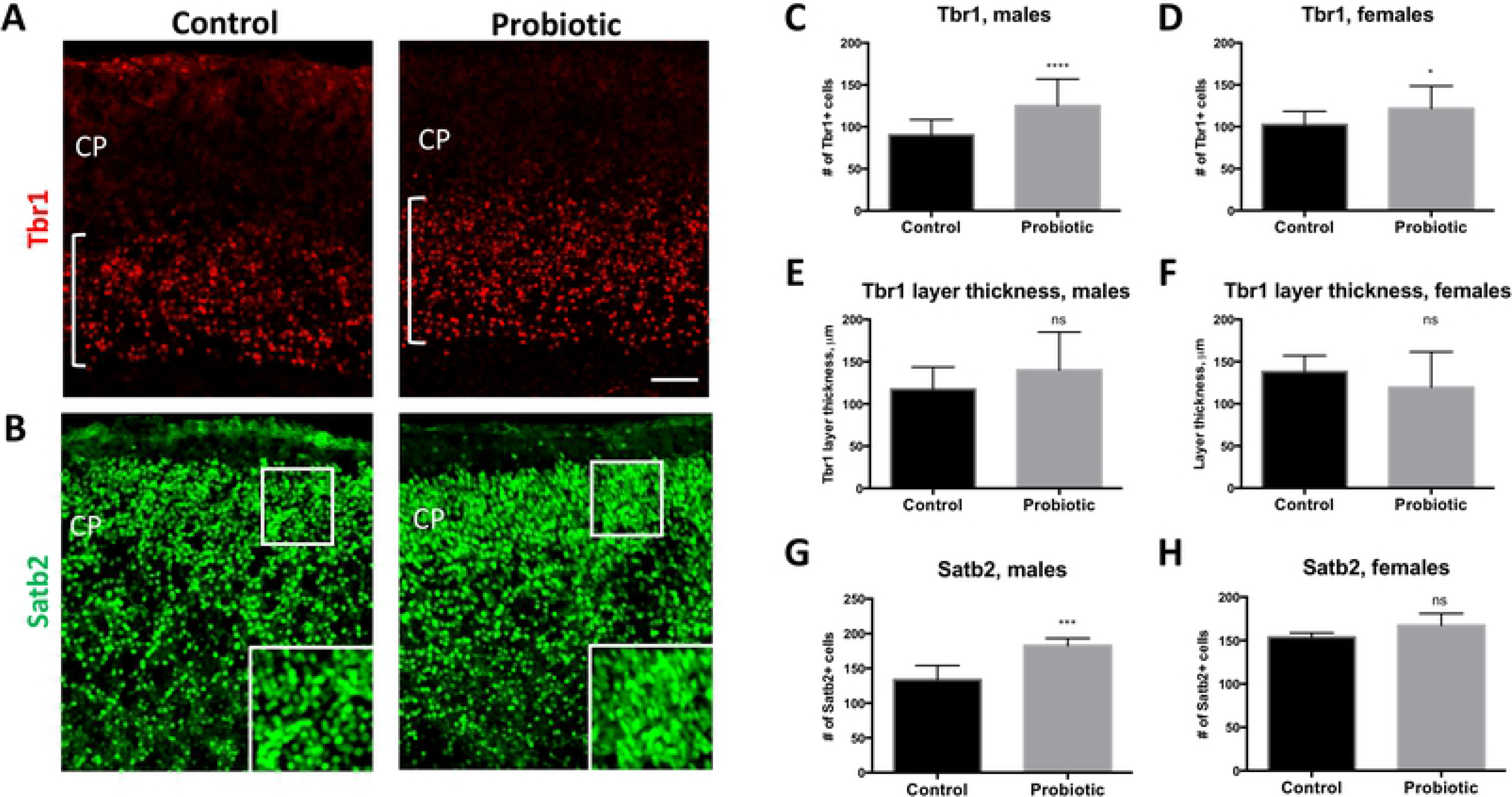
Changes in cortical neuronal marker expression in P1 pups induced by maternal intake of probiotics during pregnancy. (**A**) Expression of Tbr1 is increased in the cortices of P1 pups exposed to *L. lactis*, compared to control pups. White brackets indicate Tbr1 layer thickness. (**B**) Density of Satb2-expressing cells in the upper layer of the cortical wall is increased in P1 *L. lactis*-exposed compared to control pups. Insets show higher magnification images of areas designated by white boxes. (**C-D**) Density of Tbr1-expressing cells is increased in male (**C**; p < 0.0001 by Two-tailed Student’s t-test; control n = 26; *L. lactis* n = 13) and female (**D**; p = 0.05 by Two-tailed Student’s t-test; control n = 10; *L. lactis* n = 18) probiotic-exposed P1 pups. (**E-F**) Tbr1 layer thickness is not significantly increased in probiotic-exposed compared to control P1 pups. (**G-H**) Density of Satb2-expressing cells is increased in the upper cortical layers (II-IV) of probiotic-exposed P1 male pups (**G**; p = 0.0006 by Two-tailed Student’s t-test; control n = 7; *L. lactis* n = 5), but not female pups (**E**; p = 0.16 by Two-tailed Student’s t-test; control n = 5; *L. lactis* n = 5) compared to control groups. Scale bar in (**A**): 80 μm. *CP* - cortical plate.

Cortical pyramidal neurons and glia ultimately arise from neural progenitor cells (NPCs) that reside in the ventricular zone ^[15]^. We therefore sought to examine whether the proliferative capacity of early postnatal NPCs may be affected by exposure to probiotic. We find that the density of mitotic NPCs expressing phosphorylated histone H3 (PH3), a marker of mitosis, is increased in the ventricular zone of probiotics-exposed female (Figure 6A, C; p = 0.0009 by Student’s t-test), but not male pups, compared to P1 control female and male pups. Together, these results suggest that proliferative properties of postnatal cortical NPCs, as well as genesis and placement of cortical neurons within the cortical plate, are modulated by maternal exposure to probiotics.

**Figure 6.**
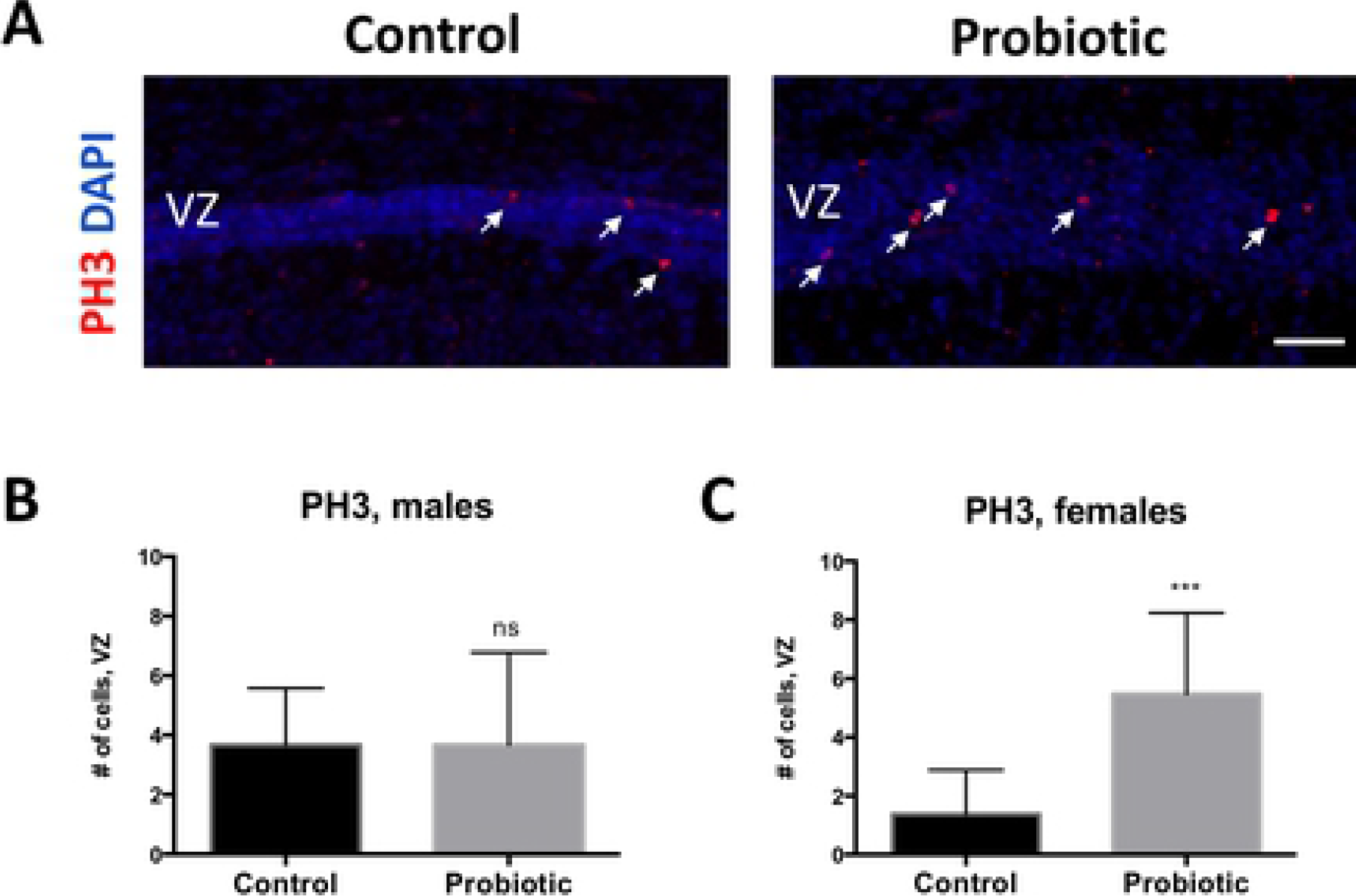
Expression of mitotic marker PH3 is increased in probiotics-exposed female P1 pups. (**A**) Larger numbers of PH3-expressing cells (white arrows) are observed in the cortical ventricular zone (VZ) of *L. lactis*-exposed versus control P1 pups. (**B-C**) Increase in PH3-expressing cells is significant in female pups (**C**; p = 0.0009 by Two-tailed Student’s t-test; control n = 8; *L. lactis* n = 15), but not in male pups (**B**) (p > 0.9 by Two-tailed Student’s t-test; control n = 18; *L. lactis* n = 15), compared to control groups. Scale bar in A: 60 μm.

## Discussion

As observed in the present study and in previous studies, prenatal probiotic treatment may have long-lasting effects on behavior. Although the supposed sterility of fetal gut is now being disputed, previous studies have repeatedly shown that colonization of the neonatal microbiome can be altered via maternal stress and/or diet ^[17]^. For example, the gut microbiomes of humans or rodents exposed to prenatal stress tend towards less diversity and numbers of lactic acid-producing strains compared to the microbiomes of unstressed offspring ^[2]^. Additionally, developmental exposure to *Bifidus* and *Lactobacillus* strains have been shown to impact gut microbiome organization and immune responses, and subsequent stress reactivity in animal models ^[4, 18]^. However, the above studies did not report the role of sex in modulation of stress via microflora interventions. Several groups have observed that maternal separation increased anxiety-like behaviors in the offspring, and probiotics treatment reversed stress effects in male rodents, but not female rodents ^[19, 20]^. Emerging evidence also indicates that sex influences microbiome colonization and alters neurotransmitter metabolism into adulthood ^[21]^.

Mice in the present study were not exposed to stress in utero, and both females and males exhibited similar anxiety-like behaviors in the open field test. However, probiotics treatment significantly increased light zone activity in the light-dark box in females only. A female-only effect of probiotics was also observed in the cue associated fear conditioning, where probiotics-treated females exhibited significantly less freezing time after a fear-associated tone than the control females. Interestingly, in this test the control females’ freezing was significantly higher than control males.

It is unclear how probiotics are differentially affecting female anxiety-like behaviors. Maternal care may differ according to sex, and interestingly dams who received probiotics exhibited higher levels of plasma oxytocin, a reputed ‘bonding’ hormone. Oxytocin enhancement of nurturing behavior may increase female offspring resiliency even in the absence of overt prenatal stress, while additional nurturing for unstressed males may confer no additional benefit. Alternatively, it may be possible that the measurement parameters of fear conditioning and light-dark box are more sensitive to sex differences than the open field test, which is more prone to variability across testing environments and time periods. Further research is needed to understand how *in utero* probiotics exposure inhibits anxiety-like behaviors, in terms of maternal versus offspring stress responses, quality of maternal care, and offspring sex.

Changes in the expression patterns of cortical vasculature, neuronal and proliferation markers in probiotic-treated mice suggest that temporal progression of cortical layer development and vascularization may be modulated by the treatment. Appropriate development of cortical architecture is dependent on a signaling sequence of growth factors, vascular integration and cross-talk between cortical layer regions. In this study, P1 pup cortices were examined for changes in the cortical layer markers, Tbr1 and Satb2, a marker of mitosis, PH3, and a marker of angiogenesis, PECAM1. Pronounced differences in cortical layer anatomy were observed in probiotics’ treated mice. Specifically, probiotics treatment induced increases in layer VI and layers II-V neuronal marker expression. In addition, vascularization in the upper layers of the cerebral wall, measured by blood vessel density, as well as average blood vessel area, were also increased in response to probiotic exposure. Taken together these results suggest that patterning of cortical structural organization in the fetus may be sensitive to maternal dietary intake of probiotics.

The immune systems of probiotic-treated neonates may be better primed *in utero* to respond to pre- and post-birth microbial exposure compared to untreated pups. The developing brain is highly sensitive to maternal immune signaling, as indicated by disrupted cortical expression patterns after maternal infections. Kim et al recently reported that mouse dam injection of synthetic double-stranded RNA, mimicking viral infection, or gavage of segmented filamentous bacteria decreased cortical expression of Satb2 in offspring, and this effect was coupled with anxiogenic behavior in an open field and social approach test ^[22]^. Furthermore, IL-17 production and gastrointestinal Th-17 cell activation was shown to be the mediator of Satb2 loss of expression. A marker of angiogenesis, CD31 (PECAM1), examined in the present study, also acts as a receptor for leukocyte activation and thus its upregulation in probiotic-treated brains may be part of an immune signaling cascade that links the gut to rate of cortical development ^[23]^. Increase in CD31 expression may also underlie the observed increase of blood vessels after probiotic supplementation.

Radial glia populations that give rise to progenitors are mainly localized to regions adjacent to brain capillaries, and rates of proliferation are influenced by signaling factors from the periphery. Along with the increased density of cortical upper layer blood vessels seen in probiotic brains, proliferation of cortical progenitors, as indicated by PH3, was also upregulated in female probiotic pups compared to control pups, and a trend effect was observed in male pups. In mice, cortical proliferation occurs during mid-gestation and subsides during early postnatal development - a temporal stage approximately equivalent to middle of the second trimester in humans. It is unknown if higher levels of PH3 expression during this time period, indicative of increased NPC proliferation and thus gliogenesis, is associated with improved resilience to stress. However, previous studies reported that decreased expression of PH3 in the developing cortex leads to anxiety-like behavior in male, but not female mice exposed to human antibodies ^[24]^. Further studies are required to delineate sex-specific pathways regulating NPC proliferation and subsequent behavior.

In conclusion, we have shown that probiotic treatment during the latter half of gestation alters cortical cytoarchitecture in P1 offspring, suggesting that changes in cortical neuronal layer and vasculature protein expression at early postnatal stages may be associated with behavioral outcomes later in life. Probiotics supplementation abrogated select anxiety-like behaviors, and improved emotional learning in females. It is presently unknown how observed changes in cortical cellular marker expression patterns may have contributed to altered behavior in probiotics-treated females, or whether changes in nurturing due to raised oxytocin levels may also have influenced sex-specific behavior differences. Further studies are needed to address the questions above and more clearly delineate the role of probiotics for human brain development.

## Acknowledgements

We thank Dr. Steven Zeisel for his input in the design of this study; Dr. David Horita for critically evaluating this manuscript; and Dr. Steven Oreña for technical assistance with oxytocin assays.

## Author contributions

N.S. and E.P. designed the study, conducted the experiments, analyzed the data, prepared the figures and wrote the manuscript; W.B.F. and C.A.M. conducted the experiments and analyzed the data; J.H. designed the study and wrote the manuscript; E.S.M. designed the study and wrote the manuscript.

## Supplementary Figure Legends

**Figure S1.**
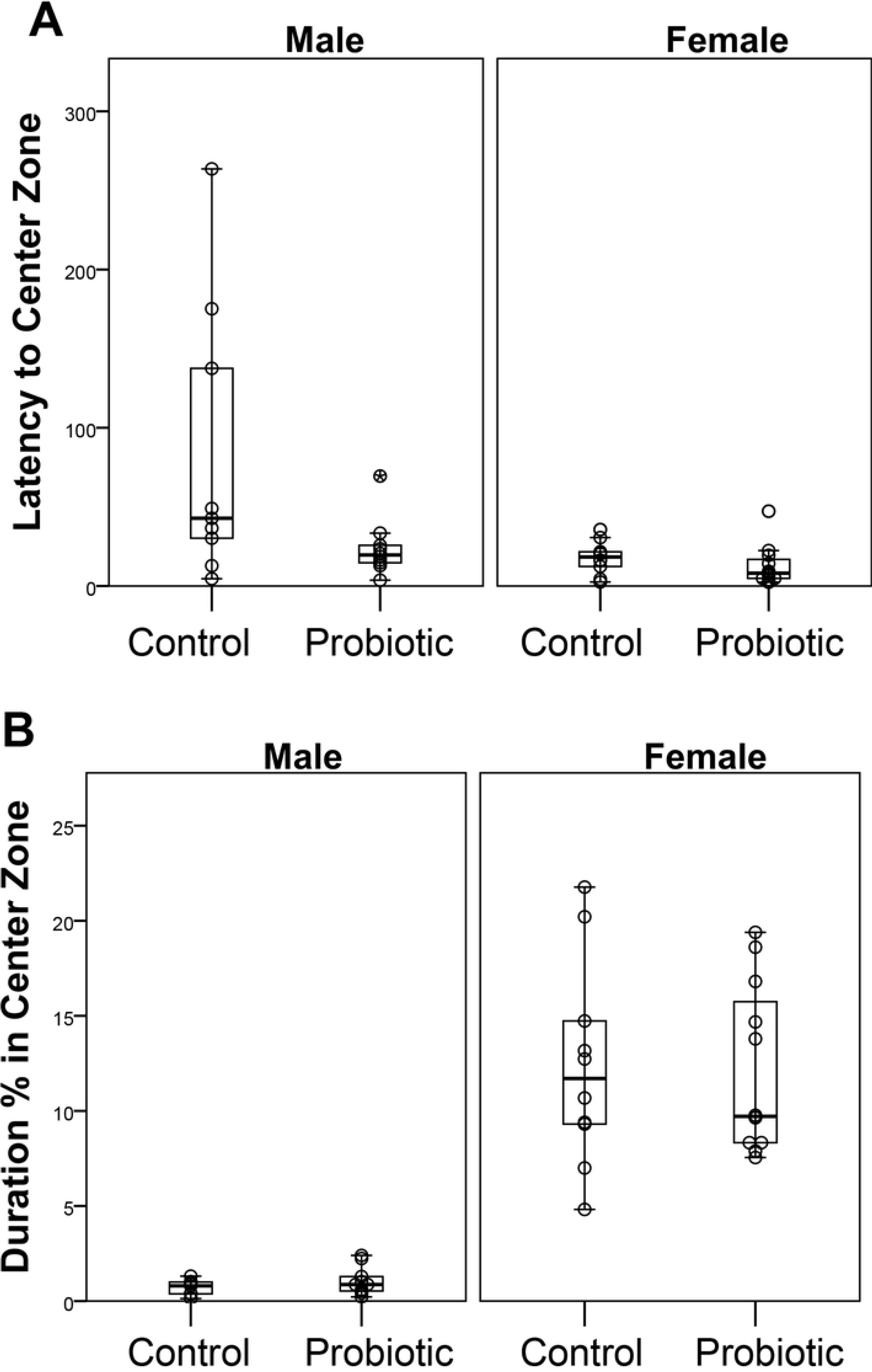
Maternal probiotic supplementation (n = 22) did not have a significant effect on open field behavior, neither on the latency to enter the center zone compared to control mice (n = 18) (**A**) nor in the time spent in the center (**B**). We did observe a sex effect, where female mice (n = 22) spend significantly more time in the center than male mice (n = 18) (**B**).

**Figure S2.**
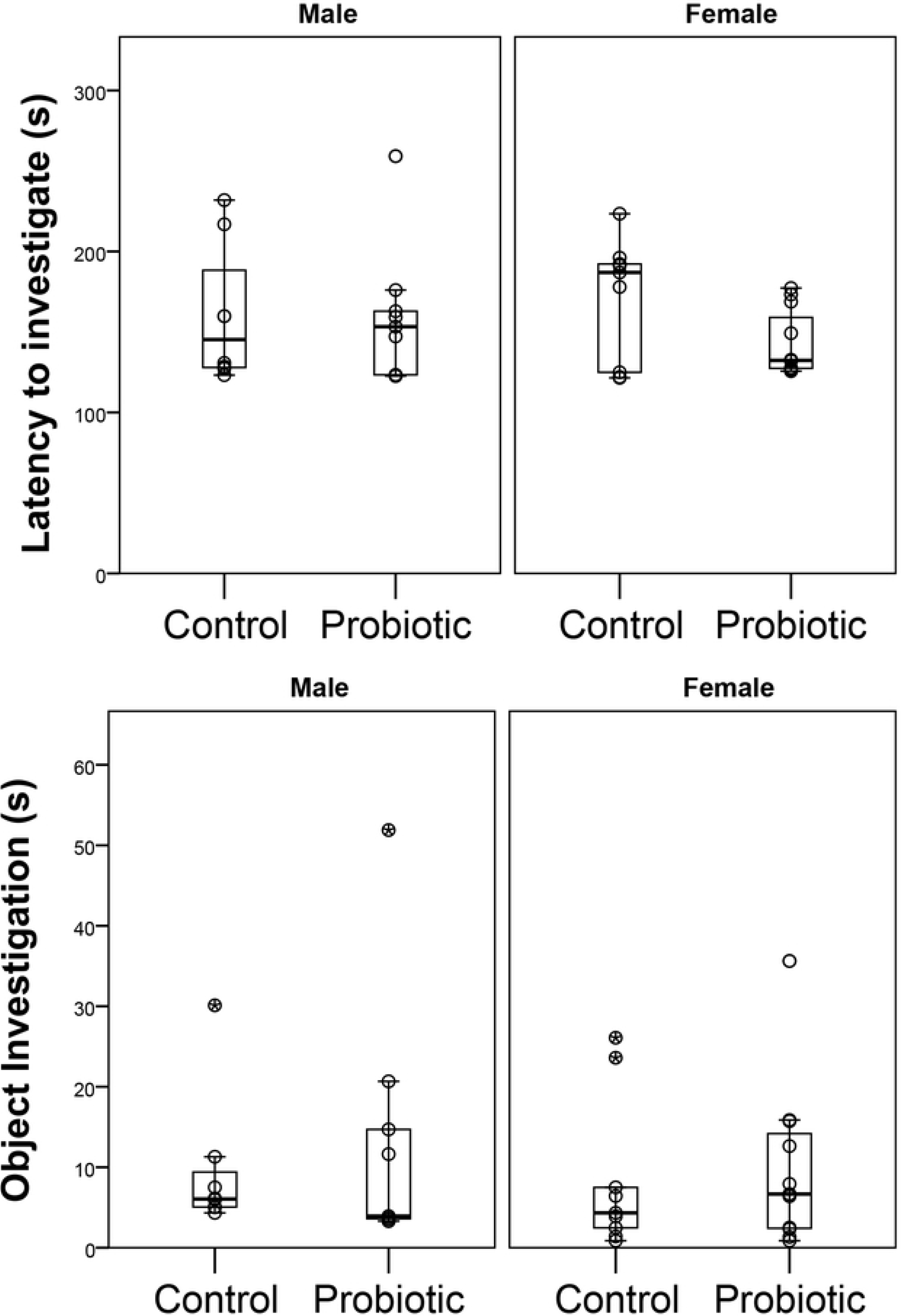
Maternal probiotic supplementation (n = 20) did not have a significant effect on novel object investigation, neither in the latency to approach the object compared to the control mice (n = 17) (**A**), nor in the time spent investigating the object (**B**).

**Figure S3.**
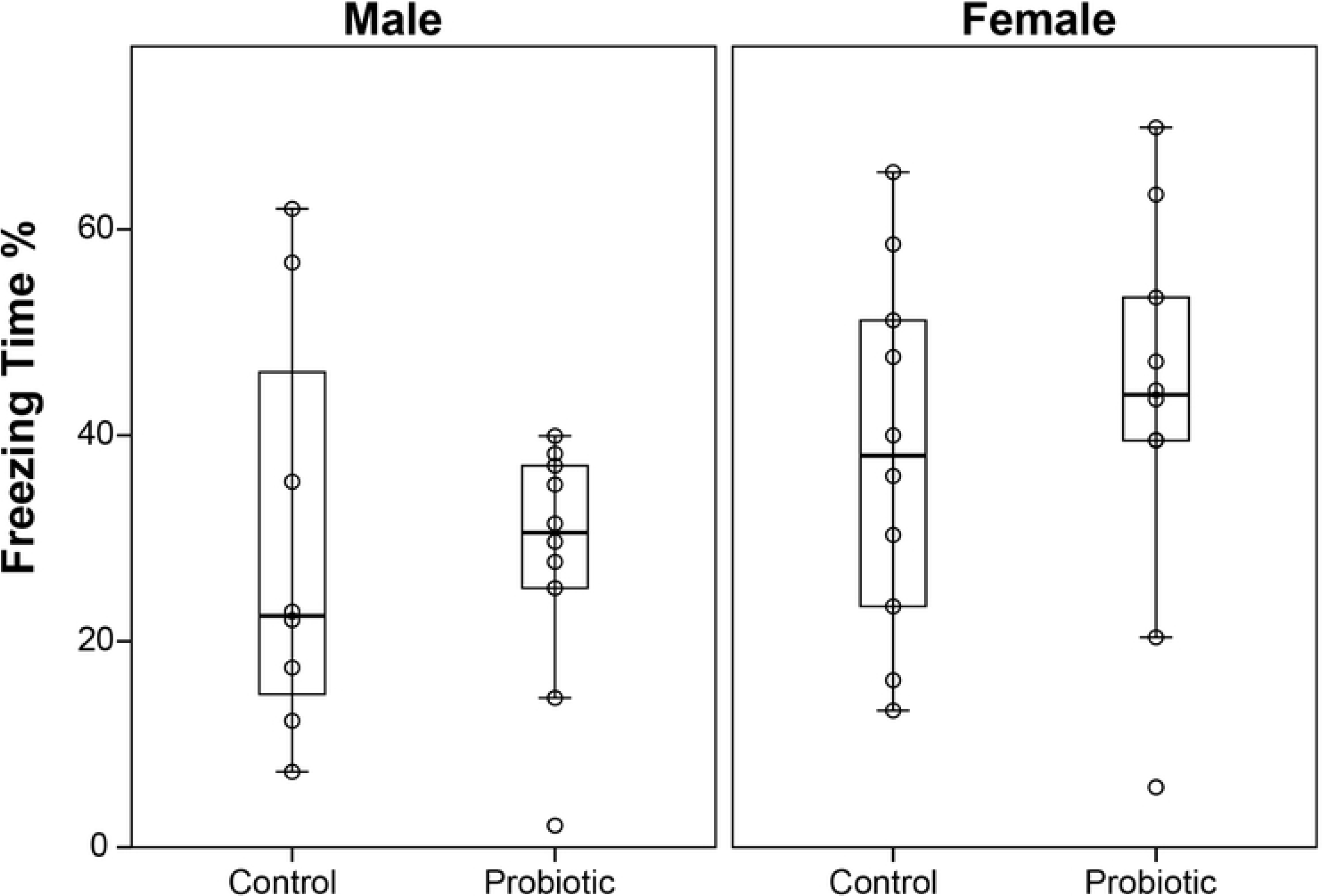
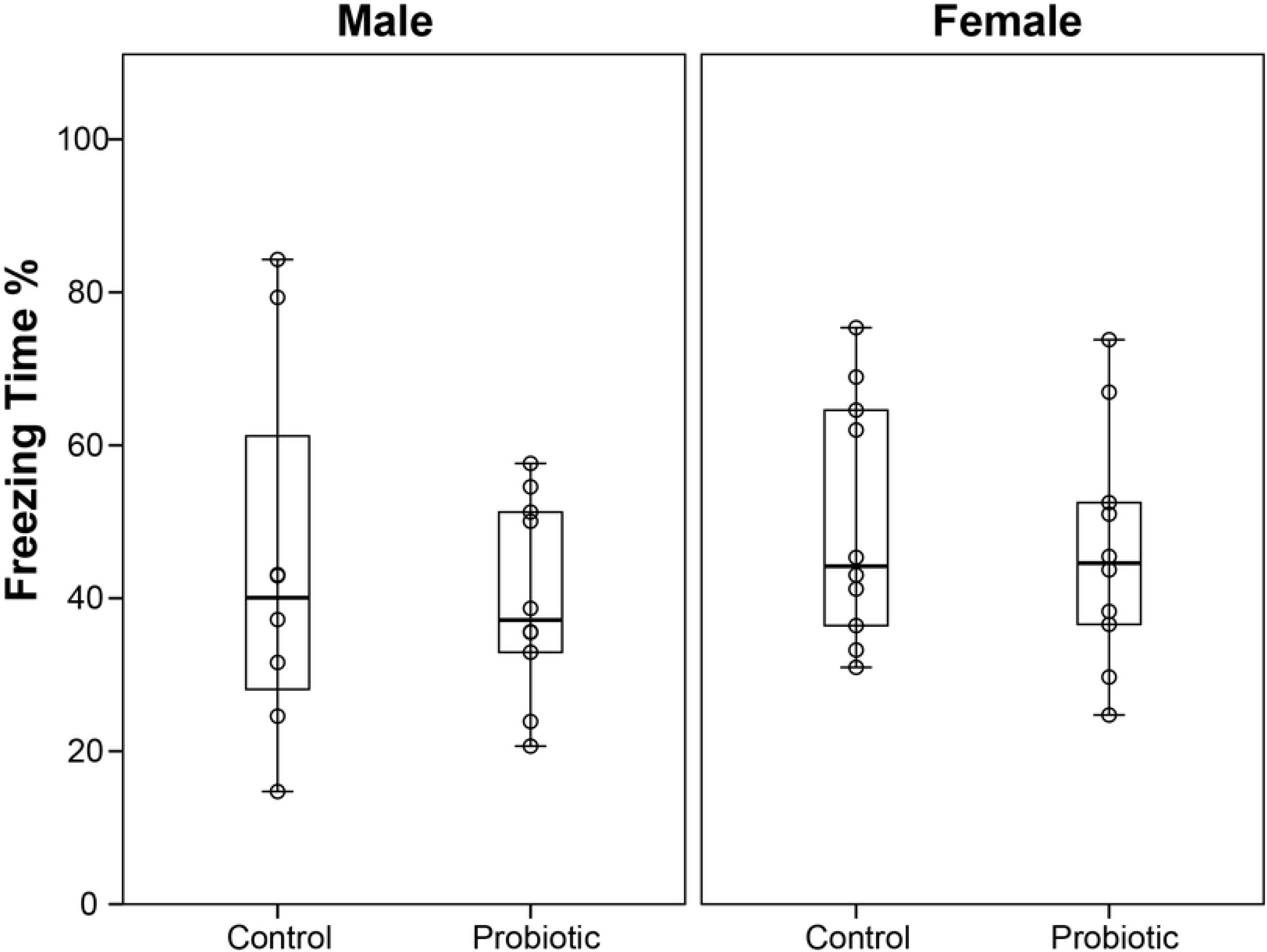
Maternal probiotic supplementation did not have an effect in contextual learning. (**A**) Female mice showed higher levels of freezing in the first minute of contextual learning but there was no treatment effect. (**B**) Probiotic exposure had no effect on average freezing time during the 5-min session showed no effect of treatment or sex.

**Figure S4.**
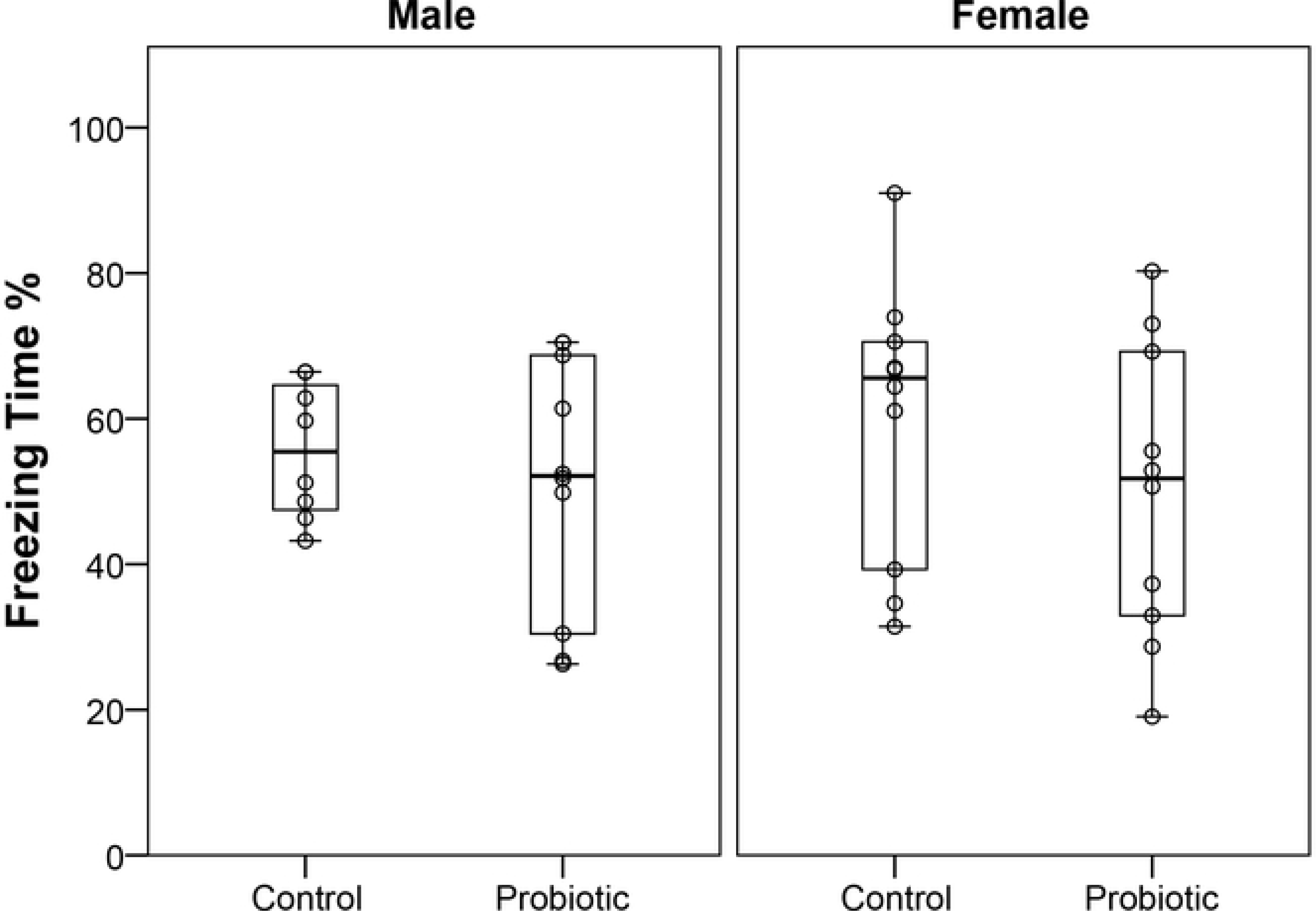
Cue associated learning. Probiotic exposure did not have an effect on the first minute response to tone in either male (control n = 8 and probiotic n = 10) or female mice (control n = 10 and probiotic n = 10).

## References

1. Wang, Y., et al., Maternal dietary intake of choline in mice regulates development of the cerebral cortex in the offspring. Faseb j, 2016. 30 (4): p. 1566–78.

2. Diaz Heijtz, R., et al., Normal gut microbiota modulates brain development and behavior. Proc Natl Acad Sci U S A, 2011. 108(7): p. 3047–52.

3. Hoban, A.E., et al., Microbial regulation of microRNA expression in the amygdala and prefrontal cortex. Microbiome, 2017. 5(1): p. 102.

4. Dinan, T.G. and J.F. Cryan, Brain-Gut-Microbiota Axis and Mental Health. Psychosom Med, 2017. 79(8): p. 920–926.

5. Hoban, A.E., et al., Regulation of prefrontal cortex myelination by the microbiota. Transl Psychiatry, 2016. 6: p. e774.

6. Luczynski, P., et al., Growing up in a Bubble: Using Germ-Free Animals to Assess the Influence of the Gut Microbiota on Brain and Behavior. Int J Neuropsychopharmacol, 2016. 19(8).

7. Savignac, H.M., et al., Bifidobacteria modulate cognitive processes in an anxious mouse strain. Behav Brain Res, 2015. 287: p. 59–72.

8. Ait-Belgnaoui, A., et al., Probiotic gut effect prevents the chronic psychological stress-induced brain activity abnormality in mice. Neurogastroenterol Motil, 2014. 26(4): p. 510–20.

9. Liu, Y.W., et al., Psychotropic effects of Lactobacillus plantarum PS128 in early life-stressed and naive adult mice. Brain Res, 2016. 1631: p. 1–12.

10. Bharwani, A., et al., Oral treatment with Lactobacillus rhamnosus attenuates behavioural deficits and immune changes in chronic social stress. BMC Med, 2017. 15(1): p. 7.

11. Bercik, P., et al., The anxiolytic effect of Bifidobacterium longum NCC3001 involves vagal pathways for gut-brain communication. Neurogastroenterol Motil, 2011. 23(12): p. 1132–9.

12. Varian, B.J., et al., Microbial lysate upregulates host oxytocin. Brain Behav Immun, 2017. 61: p. 36–49.

13. Yoshihara, C., M. Numan, and K.O. Kuroda, Oxytocin and Parental Behaviors. Curr Top Behav Neurosci, 2018. 35: p. 119–153.

14. Braniste, V., et al., The gut microbiota influences blood-brain barrier permeability in mice. Sci Transl Med, 2014. 6(263): p. 263ra158.

15. Hevner, R.F., From radial glia to pyramidal-projection neuron: transcription factor cascades in cerebral cortex development. Mol Neurobiol, 2006. 33(1): p. 33–50.

16. Agirman, G., L. Broix, and L. Nguyen, Cerebral cortex development: an outside-in perspective. FEBS Lett, 2017. 591(24): p. 3978–3992.

17. Zijlmans, M.A., et al., Maternal prenatal stress is associated with the infant intestinal microbiota. Psychoneuroendocrinology, 2015. 53: p. 233–45.

18. Cowan, C.S., B.L. Callaghan, and R. Richardson, The effects of a probiotic formulation (Lactobacillus rhamnosus and L. helveticus) on developmental trajectories of emotional learning in stressed infant rats. Transl Psychiatry, 2016. 6(5): p. e823.

19. Jasarevic, E., et al., Stress during pregnancy alters temporal and spatial dynamics of the maternal and offspring microbiome in a sex-specific manner. Sci Rep, 2017. 7: p. 44182.

20. McVey Neufeld, K.A., et al., Neurobehavioural effects of Lactobacillus rhamnosus GG alone and in combination with prebiotics polydextrose and galactooligosaccharide in male rats exposed to early-life stress. Nutr Neurosci, 2017: p. 1–10.

21. Clarke, G., et al., The microbiome-gut-brain axis during early life regulates the hippocampal serotonergic system in a sex-dependent manner. Mol Psychiatry, 2013. 18(6): p. 666–73.

22. Kim, J.A., et al., Anti-Inflammatory Effects of a Mixture of Lactic Acid Bacteria and Sodium Butyrate in Atopic Dermatitis Murine Model. J Med Food, 2018. 21(7): p. 716–725.

23. Qing, Z., et al., Inhibition of antigen-specific T cell trafficking into the central nervous system via blocking PECAM1/CD31 molecule. J Neuropathol Exp Neurol, 2001. 60(8): p. 798–807.

24. Brimberg, L., et al., Caspr2-reactive antibody cloned from a mother of an ASD child mediates an ASD-like phenotype in mice. Mol Psychiatry, 2016. 21(12): p. 1663–1671.

25. Encinas, J.M., A. Vaahtokari, and G. Enikolopov, Fluoxetine targets early progenitor cells in the adult brain. Proc Natl Acad Sci U S A, 2006. 103(21): p. 8233–8.

26. Pjetri, E. and S.H. Zeisel, Deletion of one allele of Mthfd1 (methylenetetrahydrofolate dehydrogenase 1) impairs learning in mice. Behav Brain Res, 2017. 332: p. 71–74.

27. Hall, C.S. Emotional behavior in the rat. I. Defecation and urination as measures of individual differences in emotionality. 1934. 18, 385–403.

28. Misslin, R. and P. Ropartz, Responses in Mice to a Novel Object. Behaviour, 1981. 78(3/4): p. 169–177.

29. van Gaalen, M.M. and T. Steckler, Behavioural analysis of four mouse strains in an anxiety test battery. Behav Brain Res, 2000. 115(1): p. 95–106.

30. Crawley, J. and F.K. Goodwin, Preliminary report of a simple animal behavior model for the anxiolytic effects of benzodiazepines. Pharmacol Biochem Behav, 1980. 13(2): p. 167–70.

31. Huang, H.S., et al., Behavioral deficits in an Angelman syndrome model: effects of genetic background and age. Behav Brain Res, 2013. 243: p. 79–90.

